# Cutaneous T cell lymphoma atlas reveals malignant Th2 cells supported by a B cell-rich microenvironment

**DOI:** 10.1101/2023.11.06.565474

**Authors:** Ruoyan Li, Johanna Strobl, Elizabeth F.M. Poyner, Fereshteh Torabi, Pasha Mazin, Nana-Jane Chipampe, Emily Stephenson, Louis Gardner, Bayanne Olabi, Rowen Coulthard, Rachel A. Botting, Nina Zila, Elena Prigmore, Nusayhah Gopee, Justin Engelbert, Issac Goh, Hon Man Chan, Harriet Johnson, Jasmine Ellis, Victoria Rowe, Win Tun, Gary Reynolds, April Rose Foster, Laure Gambardella, Elena Winheim, Chloe Admane, Benjamin Rumney, Lloyd Steele, Laura Jardine, Julia Nenonen, Keir Pickard, Jennifer Lumley, Philip Hampton, Simeng Hu, Fengjie Liu, Xiangjun Liu, David Horsfall, Daniela Basurto-Lozada, Louise Grimble, Chris M. Bacon, Sophie Weatherhead, Hanna Brauner, Yang Wang, Fan Bai, Nick J. Reynolds, Judith E. Allen, Constanze Jonak, Patrick M. Brunner, Sarah A. Teichmann, Muzlifah Haniffa

## Abstract

Cutaneous T-cell lymphoma (CTCL) is a potentially fatal clonal malignancy of T cells primarily affecting the skin. The most common form of CTCL, mycosis fungoides (MF), can be difficult to diagnose resulting in treatment delay. The pathogenesis of CTCL is not fully understood due to limited data from patient studies. We performed single-cell RNA sequencing and spatial transcriptomics profiling of skin from patients with MF-type CTCL, and an integrated comparative analysis with human skin cell atlas datasets from healthy skin, atopic dermatitis and psoriasis. We reveal the co-optation of Th2-immune gene programmes by malignant CTCL cells and modelling of the tumour microenvironment to support their survival. We identify MHC-II^+^ fibroblast subsets reminiscent of lymph node T-zone reticular cells and monocyte-derived dendritic cells that can maintain Th2-like tumour cells. CTCL Th2-like tumour cells are spatially associated with B cells, forming aggregates reminiscent of tertiary lymphoid structures which are more prominent with progressive disease. Finally, we validated the enrichment of B cells in CTCL skin infiltrates and its association with disease progression across three independent patient cohorts. Our findings provide diagnostic aids, potential biomarkers for disease staging and therapeutic strategies for CTCL.

## Introduction

Cutaneous T-cell lymphoma (CTCL) is a rare disease^1^ and a subgroup of non-Hodgkin lymphomas, with mycosis fungoides (MF) being the most common type with an incidence of 5.42 per million in the United States^2^. Early-stage MF (stages I-IIA^3^) typically presents in the skin with patches and plaques, which can be mistaken for benign inflammatory conditions such as atopic dermatitis (AD) and psoriasis, posing a challenge for clinical and histological diagnosis^4,5^. Although indolent in the majority, one third of patients with MF can progress to advanced-stage disease (≥ IIB) with low overall survival^6,7^. Malignant T cells in advanced-stage MF are typically central memory-like CD4^+^ clones characterised by high inter-donor variability^8^ and high tumour mutational burden^9^, but very little is known about their molecular characteristics or metabolic activity.

Identification of reliable diagnostic hallmarks for CTCL across patients has been limited by its non-specific histopathological features, as well as the heterogeneity and proposed plasticity of malignant T cells^10^. Current diagnosis is mainly based on correlation of clinical and non-specific histopathological features, including T cell epidermotropism, band-like dermal infiltrate and fibrosis of the papillary dermis^11^, all of which can also be observed in benign inflammatory skin conditions. As such, early CTCL has been termed ‘the great imitator’^12^. Research into CTCL has traditionally focused on tumour cells in peripheral blood from patients with advanced disease^13,14^. More recent studies on skin lesions from CTCL patients have largely been on small patient numbers^15^ and primarily focused on tumour cells^16^. Molecular characterisation of malignant T cells has led to non-curative treatment options for advanced CTCL, including a monoclonal antibody directed against CCR4 (mogamulizumab)^17^ and a CD30 antibody-drug conjugate (brentuximab vedotin)^18^. A subset of patients has also shown to respond to anti-PD-1 immunotherapy^19,20^.

In this study, we aimed to achieve a holistic understanding of tumour cells and their microenvironment in lesional skin from CTCL patients and integrate with data from previous studies. We performed single-cell RNA sequencing (scRNA-seq), T cell receptor (TCR) sequencing and spatial transcriptomics on skin biopsies from early and advanced stage MF-type CTCL patients, and performed a comparative analysis with single-cell and bulk RNA-seq datasets of CTCL, healthy skin, AD and psoriasis^16,21-24^. Our study identified a predominance of Th2-like malignant T cells in CTCL tumours which were likely supported by MHC-II^+^ fibroblasts and dendritic cells within the TME. In addition, we demonstrate an association of malignant Th2-like cells with B cell aggregations and with progressive disease. Finally, we demonstrate the formation of tertiary lymphoid structures in CTCL lesional skin. Our findings provide diagnostic aids, potential biomarkers for disease staging and therapeutic strategies for CTCL.

## Results

### Cellular and molecular composition of CTCL and the tumour microenvironment

We sampled epidermal and dermal lesional skin biopsies from eight (early and advanced-stage) CTCL patients with MF (Supplementary Table 1) and performed droplet-based 5’ scRNA-seq with TCR enrichment (10x Chromium platform) of all live cells from the CD45^+^CD8^+^, CD45^+^CD8^-^ and CD45^-^ fractions following FACS isolation (Fig. 1a, Extended Data Fig. 1a), as well as spatial transcriptomics analysis of tissue sections (10x Visium). Following quality control and doublet removal (see Methods), we captured ∼280,000 single cells which could be broadly categorised into 11 cell types based on the expression of canonical marker genes (Fig. 1b, Extended Data Fig. 1b,c). A representative view of CTCL tumour microenvironment (TME) is shown by multi-colour immunofluorescence imaging showing T cells in both epidermis and dermis, together with myeloid cells and B cells surrounding dermal blood vessels (Fig. 1c). Our scRNA-seq and spatial transcriptomics data can be explored using WebAtlas^25^ (https://collections.cellatlas.io/ctcl).

**Fig. 1.**
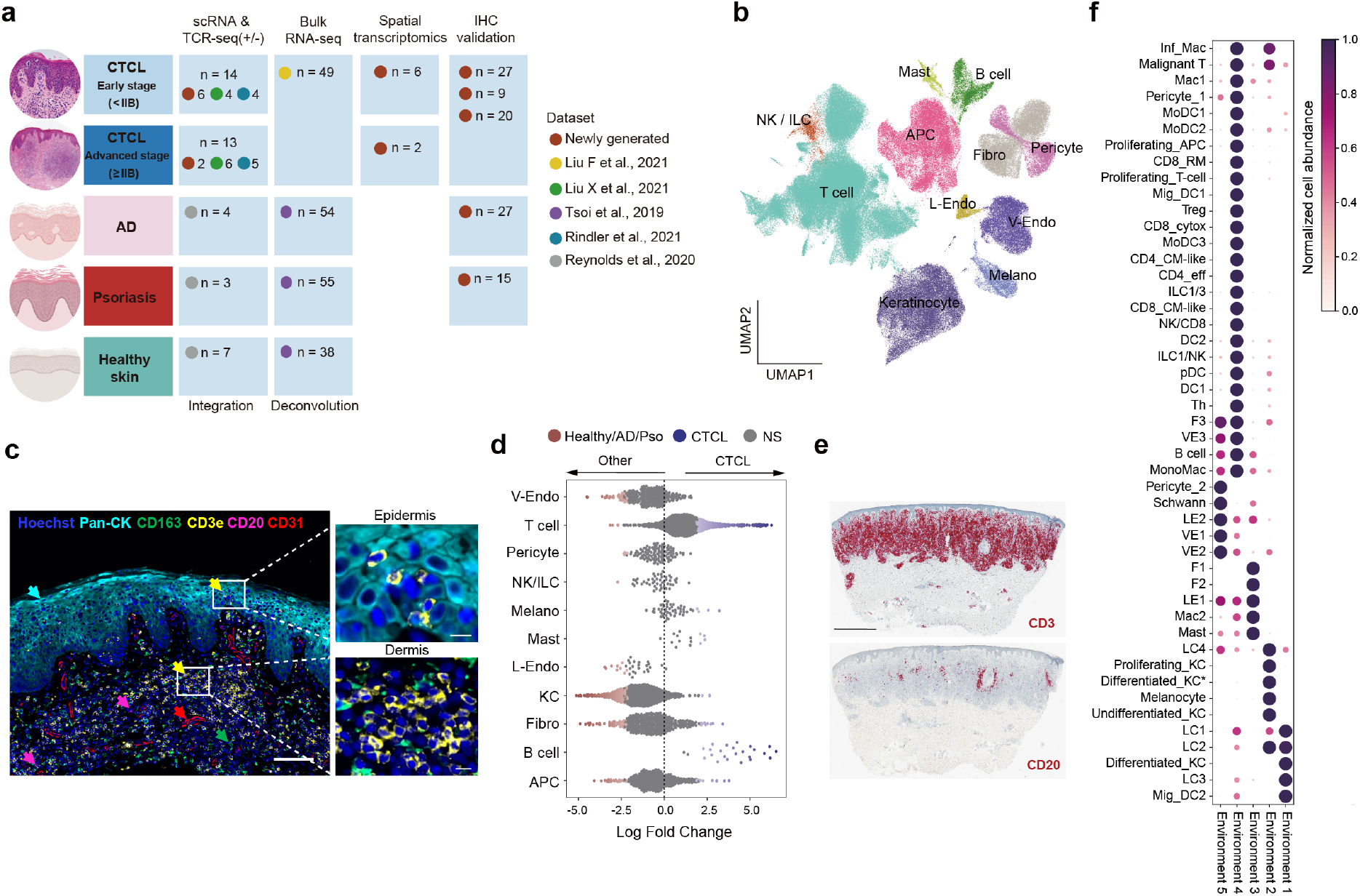
Overview of the CTCL dataset and comparisons to skin cell atlas. **a**, Detailed summary of newly generated data and integrated external datasets in this study. **b**, Overall uniform manifold approximation and projection (UMAP) showing major cell types in our CTCL dataset (NK, natural killer cell; ILC, innate lymphoid cell; APC, antigen presenting cell; Fibro, fibroblast; V-Endo, vascular endothelial cell; L-Endo, lymphatic endothelial cell; Melano, melanocyte). **c**, A multi-colour immunofluorescence image (Rarecyte) showing a representative view of CTCL TME. Scale bars: 100 μm (zoomed-out) and 10 μm (zoomed-in). **d**, Beeswarm plot of the log-transformed fold changes in abundance of cells in CTCL versus those in healthy skin, AD and psoriasis from skin cell atlas. Differential abundance neighbourhoods at FDR 10% are coloured. NS, not significant. **e**, IHC staining for CD3 and CD20 in a representative sample. Scale bar: 1 mm. **f**, Dot plot showing estimated non-negative matrix factorisation (NMF) weights of cell types across NMF factors (Environments). Shown are relative weights, normalised across factors for every cell type.

We integrated our scRNA-seq CTCL data with an existing human skin cell atlas dataset^22^, which includes healthy, AD and psoriasis skin to distinguish CTCL-specific features from common inflammatory skin disorders (Fig. 1a, Extended Data Fig. 1d). We performed label transfer from the skin cell atlas using logistic regression to annotate cell states in the integrated dataset (Extended Data Fig. 1e). Differential abundance testing using Milo^26^ was performed to interrogate differences in cellular abundance for the broad cell types seen in CTCL compared to healthy skin, AD and psoriasis (Fig. 1d). We observed an expected enrichment of T cells (Fig. 1d) but also enrichment of B cells and mast cells in CTCL (Fig. 1d). The enrichment of B cells is surprising as they are not usually present in the skin, whether in healthy, AD and psoriasis contexts^22^. We further confirmed the presence of B cells in CTCL by immunohistochemical staining (IHC; Fig. 1e).

We mapped the fine-grained annotated cell types to their spatial locations in skin tissues using cell2location^27^. Analysis revealed malignant T cells sharing a microenvironment with fibroblasts, dendritic cells and B cells in the dermis (Environment 4) as well as undifferentiated and differentiated keratinocytes, and Langerhans cells (LCs) in the epidermis (Environment 2) (Fig. 1f).

### Malignant T cells in CTCL

To distinguish malignant/tumour cells from benign infiltrating T cells in CTCL skin, we inferred large-scale chromosomal copy number variations (CNVs) based on scRNA-seq data (see Methods). As expected, malignant T cells exhibited extensive CNVs across their genomes consistent with those identified from whole-genome sequencing of the same tumour (Extended Data Fig. 2a).

In contrast to benign infiltrating T cells which clustered together across patients, malignant T cells from CTCL skin clustered separately according to patient origin (Fig. 2a). Further sub-clustering of benign T cells, ILCs and NK cells identified 12 cell subsets in CTCL, healthy skin, AD and psoriasis (Extended Data Fig. 2b,c), showing different expression features (Extended Data Fig. 2d). Differential abundance testing revealed enrichment of regulatory T cells (Treg) and CD8^+^ T cells with a cytotoxic profile (CD8_cytox) in CTCL (Extended Data Fig. 2e-g). Overall, benign lymphocytes in CTCL resemble T cells of the TME in other cutaneous squamous and melanocytic cancer types, including melanoma, with high abundance of cytotoxic CD8^+^ T cells, potentially pro-tumourigenic CD4^+^ T helper cells and Tregs^28,29^.

**Fig. 2.**
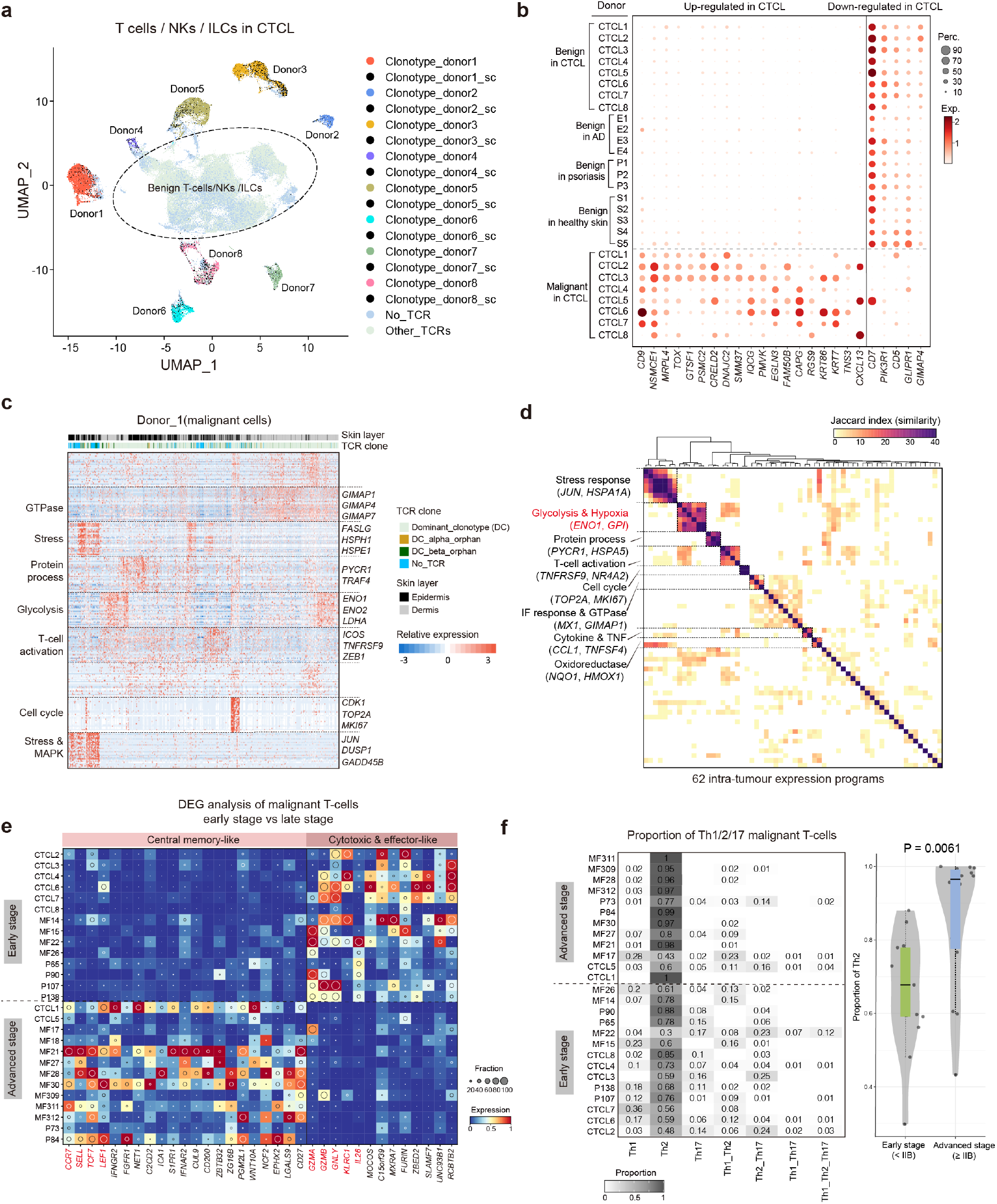
Characterisation of malignant T cells in CTCL. **a**, UMAP visualisation of T, NK and ILC cells in our CTCL dataset. Colours highlight the predominant T cell clonotypes in each patient. Black dots represent cells that share one TCR chain with the predominant T cell clonotype in a specific patient. **b**, Dot plot of DEGs that are up-regulated or down-regulated in malignant T cells compared to benign T cells from healthy, AD, psoriasis and CTCL skin. Dot colours indicate log-transformed and normalised expression values. Dot size indicates the percentage of cells expressing a given gene. **c**, Heat map showing gene expression programmes and intra-tumour expression heterogeneity among malignant T cells in a representative patient. Programme annotations and representative genes are labelled. Colour key indicates scaled expression levels. **d**, Heat map depicting shared expression meta-programmes across all patients. Jaccard index is used to measure the similarity between any two intra-tumour expression programmes. **e**, Heat map of DEGs in early or advanced stage CTCL samples. Colour represents expression level standardised between 0 and 1. The inset circle indicates the percentage of cells expressing a given gene. **f**, Left, heat map showing the proportion of putative Th1-, Th2- and Th17-like malignant cells in each CTCL patient. Patients are categorised into early or advanced stages using stage IIB as a boundary. Right, violin plot comparing the proportion of Th2-like malignant cells in early and advanced stage CTCL samples. P value is calculated using a two-sided Wilcoxon rank sum test. The lower edge, upper edge and centre of the box represent the 25th (Q1) percentile, 75th (Q3) percentile and the median, respectively. The interquartile range (IQR) is Q3 – Q1. Outliers are values beyond the whiskers (upper, Q3 + 1.5 × IQR; lower, Q1 − 1.5 × IQR).

Cross-tumour comparisons revealed differentially expressed genes (DEGs) between malignant T cell clones (Fig. 2a, Extended Data Fig. 2h,i). For example, malignant T cells from patient 4 were highly cytotoxic-like with high expression of genes such as *IFNG* and *GZMB* (Extended Data Fig. 2i). Further projection of dominant TCR clonotypes onto the UMAP revealed that malignant T cells from each patient almost exclusively harboured a single clonally expanded TCR (Fig. 2a), in keeping with published reports on CTCL tumour cells^16^. Notably, some malignant cells had lost either one or both TCR chains, consistent with known loss of T cell phenotypic identity by histopathological staining and observed in scRNA-seq profiling^21^ (Fig. 2a).

To further distinguish the molecular properties of malignant T cells from benign infiltrating T cells in CTCL, we analysed DEGs between malignant T cells and benign T cells from healthy skin, AD, psoriasis and CTCL. In total, we identified 767 upregulated and 592 downregulated DEGs in malignant T cells, including previously reported features such as *TOX* upregulation and *CD7* downregulation^30,31^ (Fig. 2b, Supplementary Table 2). 24 genes provided good discriminatory power to distinguish malignant from benign T cells (Fig. 2b). For instance, *CD9*, which encodes a cell surface glycoprotein, was upregulated in seven out of eight tumours (Fig. 2b). Interestingly, we observed high expression of *CXCL13* in malignant T cells from three tumours (Fig. 2b), suggestive of a B cell homing and recruitment role^32^.

We corroborated the upregulated DEGs observed in multiple tumours (Fig. 2b) in two published CTCL scRNA-seq datasets^16,21^ (Extended Data Fig. 3a). In addition, we validated protein expression of TOX and GTSF1, previously reported to distinguish malignant CTCL cells^33^, in lesional skin using IHC (n=13) and observed increased, but highly variable, expression of these markers (Extended Data Fig. 3b,c). Although these upregulated genes in malignant T cells show potential as biomarkers, no marker alone could identify all CTCL tumours, demonstrating the heterogeneity of CTCL and the need to profile the presence of several genes/markers for diagnostic precision.

### Metabolic gene programmes in malignant T cells are conserved across patients

To further dissect intra-tumoural malignant T cell heterogeneity in CTCL and identify features shared across all CTCL tumours, we analysed intra-tumoural co-expressed gene modules (Fig. 2c). We identified 62 intra-tumour expression programmes in total and classified eight meta-programmes (MPs) shared by subpopulations of malignant T cells in multiple tumours (Fig. 2d, Supplementary Table 3). Among these MPs, a glycolysis and hypoxia-related MP was shared by six tumours and characterised by the expression of genes such as *ENO1* and *GPI* (Fig. 2d, Supplementary Table 3). A subset of malignant cells from all donors highly expressed aerobic glycolysis pathway genes. These genes are also highly expressed by a proportion of benign T cells in CTCL. Interestingly, this metabolic feature is shared by cancer cells in hypoxic environments and circulating memory T cells, but not tissue-resident T cells^34^, and may present a therapeutic target in CTCL yet to be explored.

Malignant T cells initially exhibit epidermotropism but subsequently migrate into the dermis with CTCL severity^35^. By sampling and profiling epidermal and dermal lesions separately, we could compare gene expression between malignant T cells from both compartments to capture this process. This revealed higher expression of cell migration-related genes including *CCR7* in dermal malignant T cells, in contrast to higher expression of metabolism-related genes like *NQO1/2, FABP5* and *PRDX1* in epidermal malignant T cells (Extended Data Fig. 3d, Supplementary Table 4). This finding suggests dermal and epidermal malignant T cells possess differential migratory potential and adaptation to local microenvironmental nutrient availability.

### Malignant T cells in advanced-stage CTCL exhibit Th2-skewing and a central memory-like expression profile

We further integrated our tumour cell data with two published scRNA-seq datasets^16,21^ and focused on the differential gene expression between malignant T cells from early-stage (< IIB stage) and advanced-stage CTCL (≥ IIB stage). In early-stage samples, malignant T cells were cytotoxic and tissue resident effector-like, as reflected by higher expression of *GZMA*, *GZMB* and *GNLY*, and lower expression *CCR7*, while in advanced stage, malignant T cells expressed features of central memory cells including *SELL*, *CCR7, LEF1* and *TCF7* (Fig. 2e, Supplementary Table 5), suggesting the capacity to circulate in more advanced disease. This pattern is also observed in bulk RNA-seq data (Extended Data Fig. 3e; both *P*<0.05, two-sided Wilcoxon rank-sum test). Next, we determined the functional phenotype of malignant T cells as Th1-, Th2- and Th17-like cells based on the expression of T cell lineage transcription factors *TBX21*, *GATA3* and *RORC* respectively, and compared the proportion of Th types across disease stages. Compared to early-stage samples, a significantly higher proportion of Th2-like malignant T cells was observed in advanced-stage samples (Fig. 2f; *P*=0.0061, two-sided Wilcoxon rank-sum test), suggesting either Th2-skewing of Th1/Th17-like malignant cells or preferential survival of Th2-like malignant cells upon CTCL disease progression.

### MHC-II^+^ fibroblasts likely support malignant cells in CTCL

We next interrogated the stromal cell compartment in CTCL compared to healthy skin, AD and psoriasis. Sub-clustering and annotation of stromal and KC populations (referred to as stromal population) identified 16 cell states in CTCL, healthy, AD and psoriasis skin (Fig. 3a, Extended Data Fig. 4a,b). We next focused on differential cellular abundance in the stromal subtypes in CTCL compared to healthy skin, AD and psoriasis (Fig. 3b, Extended Data Fig. 4c). The greatest differential abundance and qualitative gene expression changes were observed in fibroblast subtypes F2 and F3, where a number of cell neighbourhoods were significantly enriched in CTCL (Fig. 3b). We performed deconvolution analysis of bulk RNA-seq datasets^23,24^ using BayesPrism^36^, which validated the enrichment of F2 (Extended Data Fig. 4d; all *P*<10^-4^, two-sided Wilcoxon rank-sum test).

**Fig. 3.**
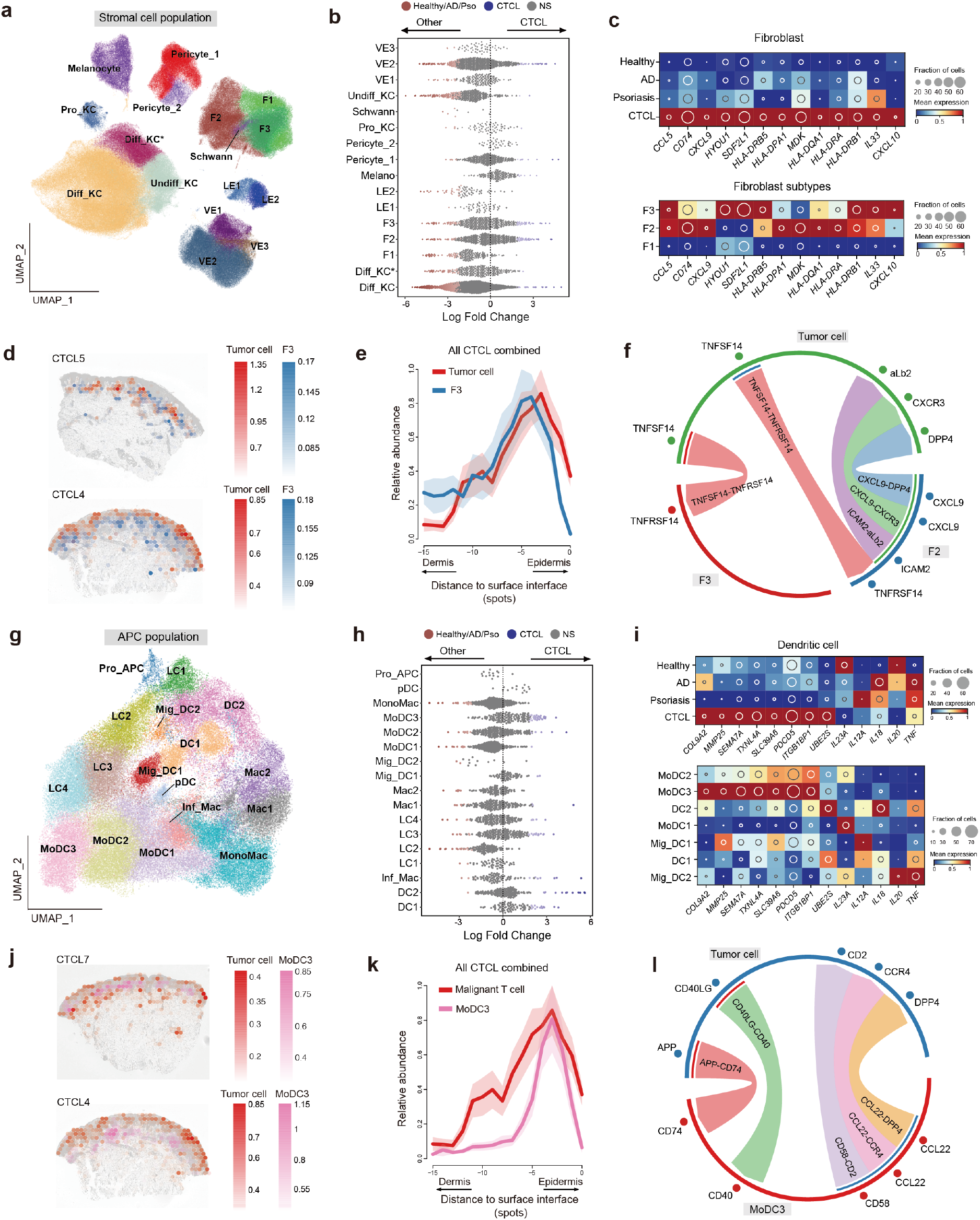
Characterisation of the stromal and APC compartments in CTCL. **a**, UMAP visualisation of the stromal population in the current dataset integrated with skin cell atlas, coloured by cell types (VE, vascular endothelial cells; KC, keratinocytes; Melano, melanocytes; LE, lymphatic endothelial cells; F, fibroblasts). **b**, Beeswarm plot of the log-transformed fold changes in abundance of stromal cell subsets in CTCL versus those in healthy skin, AD and psoriasis. Differential abundance neighbourhoods at FDR 10% are coloured. NS, not significant. **c**, Heat map showing gene expression in CTCL enriched fibroblasts. Colour represents expression level standardised between 0 and 1. The inset circle indicates the percentage of cells expressing a given gene. **d**, Spatial mapping of F3 and tumour cells in two representative samples. Estimated abundance (colour intensity) is overlaid on histology images. **e**, Curve plot showing the mean (across all samples) per-spot normalised abundance of F3 and tumour cells along the axis to skin surface. Shaded regions represent the 95% 2SD confidence intervals. **f**, Circos plot showing putative ligand-receptor interactions between fibroblasts and malignant T cells. Representative interactions are coloured. **g**, UMAP visualisation of the APC population in the current dataset integrated with skin cell atlas, coloured by cell types. **h**, Beeswarm plot of the log-transformed fold changes in abundance of stromal cell subsets in CTCL versus those in healthy skin, AD and psoriasis. Differential abundance neighbourhoods at FDR 10% are coloured. NS, not significant. **i**, Heat map showing gene expression in CTCL enriched DC. Colour represents expression level standardised between 0 and 1. The inset circle indicates the percentage of cells expressing a given gene. **j**, Spatial mapping of MoDC3 and tumour cells in two representative samples. Estimated abundance (colour intensity) is overlaid on histology images. **k**, Curve plot showing the mean (across all samples) per-spot normalised abundance of MoDC3 and tumour cells along the axis to skin surface. Shaded regions represent the 95% 2SD confidence intervals. **l**, Circos plot showing putative ligand-receptor interactions. Representative interactions are coloured.

To understand the function of CTCL-enriched fibroblasts, we used a pseudo-bulk strategy to analyse DEGs in fibroblasts (merging F1, F2 and F3) between CTCL and the other three conditions (Supplementary Table 6). Interestingly, we found that MHC-II genes (*CD74*, *HLA-DRB5* and *HLA-DPA1*) implicated in antigen-presenting potential were up-regulated in CTCL-enriched fibroblasts (Fig. 3c). Upregulation of these antigen presenting-related genes was predominant in CTCL-enriched F2 and F3 cell neighbourhoods that also expressed chemokines *CCL5*, *CXCL9* and *CXCL10*, and Th2-promoting cytokines such as *IL33* (Fig. 3c, Extended Data Fig. 4e). The F2/F3 CTCL skin fibroblasts resemble previously reported MHC-II^+^ antigen-presenting fibroblasts in several cancer types including pancreatic adenocarcinoma and breast cancer^37,38^, MHC-II^+^ lymph node T-zone reticular cells (TRCs) expressing CXCL9^39^, and tertiary lymphoid structure fibroblasts^40^. Furthermore, F2 fibroblasts transcriptionally resemble foetal skin fibroblasts^22^, lending support to previous reports on co-optation of developmental cell states in inflammatory disease^22^ and cancer^41^.

Spatial mapping revealed proximity between F2 and F3 fibroblasts with malignant T cells (Fig. 3d,e, Extended Data Fig. 4f). Due to the transcriptional resemblance between CTCL MHC-II^+^ fibroblasts and lymphoid organ TRCs and their spatial proximity to malignant T cells, we hypothesised that MHC-II^+^ fibroblasts may be interacting with and promoting malignant T cell growth in CTCL, analogous to TRCs supporting survival of naive T cells in the lymph node^42,43^. We therefore inferred intercellular communications between the two fibroblast subtypes and malignant T cells in CTCL based on putative ligand-receptor interactions. Our analysis predicted cell-cell interactions between F2/F3 fibroblasts with malignant T cells via ligands-receptors such as TNFSF14-TNFRSF14 and CXCL9-DPP4 (Fig. 3f). These observations suggest MHC-II^+^ fibroblasts likely support CTCL tumours in lesional skin. As well as containing MHC-II^+^ fibroblasts, CTCL lesional skin is also enriched with keratinocytes that have upregulated expression of *TSLP*, that is well recognised to promote Th2 microenvironment in AD^44^ (Extended Data Fig. 4g).

### APCs in CTCL TME promote T cell activation and Th2-skewing

To investigate if CTCL skin antigen presenting cells (APCs) supported malignant T cells, we performed sub-clustering of the APC population and annotated different subsets based on the skin cell atlas data^22^ (Fig. 3g, Extended Data Fig. 5a,b). Using differential abundance testing, we found that monocyte-derived dendritic cells (MoDCs), especially MoDC3, were substantially enriched in CTCL compared with other three conditions (Fig. 3h, Extended Data Fig. 5c). The enrichment of MoDC3 can also be found in bulk deconvolution (Extended Data Fig. 5d). Interestingly, CTCL-enriched DCs (MoDC3) showed higher expression of matrix metalloproteinase *MMP25*, and *CD40* and *CD58*, molecules known to activate T-cells through interactions with *CD40LG* and *CD2* respectively (Fig. 3i, Extended Data Fig. 5e). Indeed, we observed proximity between MoDC3 and malignant T cells (Fig. 3j,k, Extended Data Fig. 5f), and predicted interactions between them via the ligand-receptor pairs CD40-CD40LG and CD58-CD2 (Fig. 3l).

To identify CTCL-specific gene expression patterns, we compared CTCL APCs with their counterparts in healthy skin, AD and psoriasis (Extended Data Fig. 5g, Supplementary Table 6). CTCL derived LCs showed higher expression of costimulatory cytokine *CD70*, which may play potential roles in activation of malignant T cells (Extended Data Fig. 5g). Notably, genes related to Th1 and Th17 skewing (i.e., *IL23A* and *IL18*) were downregulated in DCs in CTCL skin (Fig. 3i), further supporting a Th2 permissive malignant T cell microenvironment (Fig. 3c, Extended Data Fig. 4g).

### CTCL lesional B cells form tertiary lymphoid structures and interact with malignant T cells

B cells, sometimes organised in lymphoid structures, have been reported in several cancer TMEs where they can prime and stimulate anti-tumour T cells and produce tumour-directed antibodies^45^. We therefore wanted to confirm our earlier observation of increased B cell abundance in CTCL skin in a larger cohort and if B cells were present as aggregates within lymphoid structures in CTCL lesional skin. First, we confirmed B cells were present in every CTCL patient in our scRNA-seq data, regardless of disease stage (Extended Data Fig. 6a). Then, we assessed if B cell enrichment is evident in a larger CTCL patient cohort. Using bulk deconvolution analysis (n=196), we confirmed significantly greater proportions of B cells present in CTCL skin samples compared to healthy skin, lesional and non-lesional AD and psoriatic skin (Fig. 4a; all *P*<10^-4^, two-sided Wilcoxon rank-sum test). We further validated the increased presence of B cells in CTCL skin biopsies using IHC by staining for CD20 and CD79a in three independent CTCL cohorts (n=56) (Fig. 4b, Extended Data Fig. 6b).

**Fig. 4.**
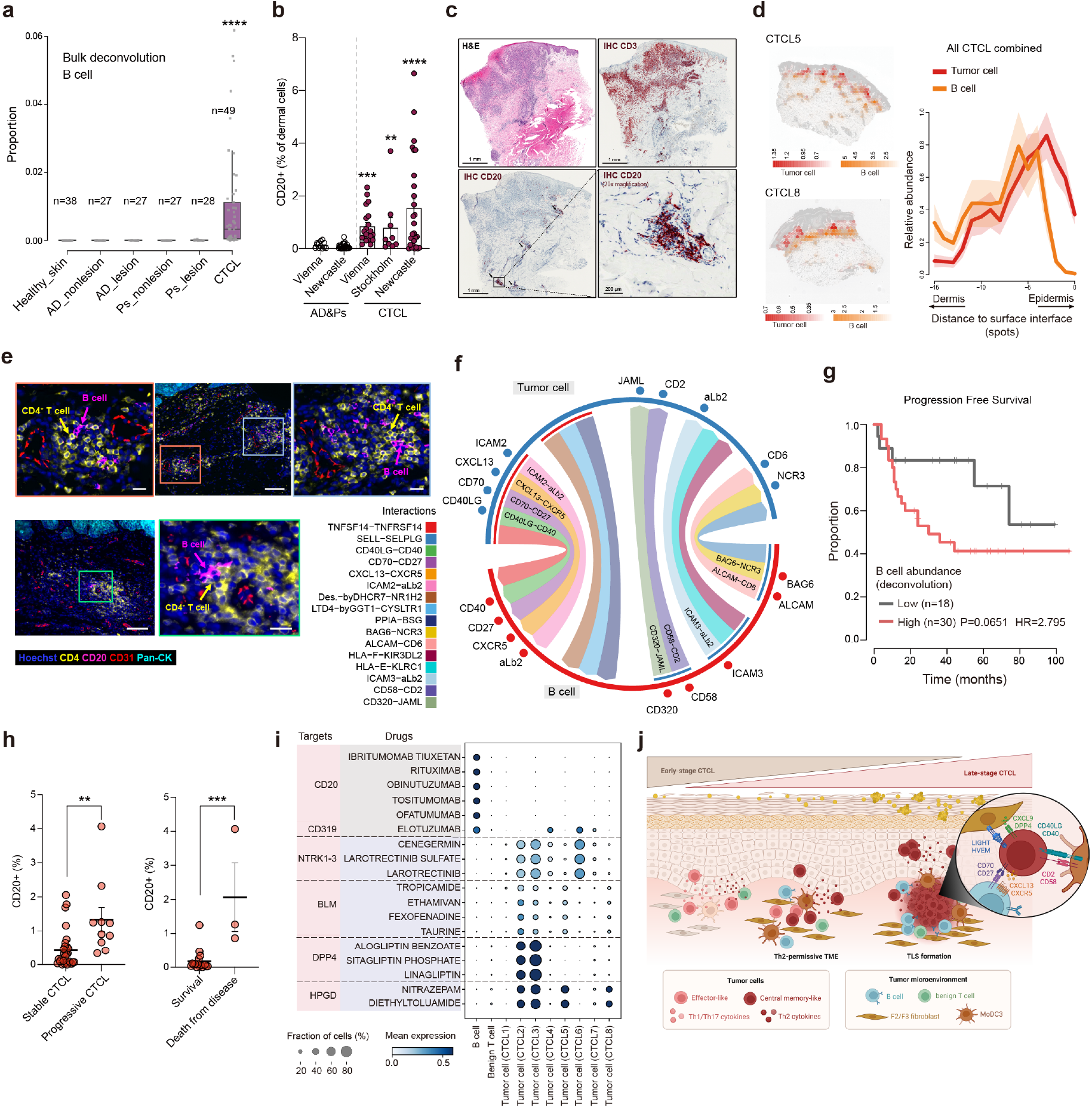
B cell enrichment in CTCL. **a**, Box plot showing deconvolution of B cells in bulk RNA-seq datasets of healthy skin, AD, psoriasis and CTCL. Numbers of samples in the categories are labelled. ****, P<0.0001. The lower edge, upper edge and centre of the box represent the 25th (Q1) percentile, 75th (Q3) percentile and the median, respectively. The interquartile range (IQR) is Q3 – Q1. Outliers are values beyond the whiskers (upper, Q3 + 1.5 × IQR; lower, Q1 − 1.5 × IQR). **b**, Bar plot showing IHC staining of CD20 in AD, psoriasis and CTCL skin samples in three independent cohorts (AD/Pso cohorts, n=27, n=15; CTCL cohorts, n=27, 9, 20). **, P<0.01, ***, P<0.001, ****, P<0.0001. Error bars show SEM. **c**, H&E image and IHC staining for CD3 and CD20 in a representative sample. The zoom-in box and arrows highlight B cells. Scale bars, 1 mm and 200 μm. **d**, Left, spatial mapping of B cells and tumour cells in two representative samples. Estimated abundance (colour intensity) is overlaid on histology images. Right, curve plot showing the mean (across all samples) per-spot normalised abundance of B cells and tumour cells along the axis to skin surface. Shades represent the 2SD intervals. **e**, Multi-colour immunofluorescence images (Rarecyte) in two representative tumours. Representative views of B cell and CD4^+^ T cell interaction are zoomed in. Scale bars: 100 μm (zoomed-out) and 20 μm (zoomed-in). **f**, Circos plot showing putative ligand-receptor interactions between B cells and malignant T cells. Representative interactions are coloured. **g**, Progression free survival probability of CTCL patients according to stratification of B cell abundance estimated by bulk deconvolution. HR, hazard ratio. **h**, IHC staining of CD20 in stable and progressive CTCL skin samples (left) and outcome (survival vs. death from disease, right). Data shown as individual values and mean percentages of CD20^+^ cells among all cells +/- SEM, n=27 (Vienna cohort), **, P<0.01, ***, P<0.001. **i**, Dot plot showing the expression of drug targets predicted by drug2cell. **j**, Schematic of the features depicting the TME of CTCL.

Notably, we found B cells formed aggregates in 55% (31/56) of CTCL IHC samples (Fig. 4c). The aggregates detected in our IHC samples were reminiscent of early tertiary lymphoid structures (TLS), which usually contain germinal centre B cells. We therefore annotated the B cell population in our CTCL dataset and identified several subsets including naive B cells and IgG-producing plasma cells (Extended Data Fig. 6c-e). Interestingly, we identified a subpopulation of germinal centre-like (GC-like) B cells highly expressing *BCL2A1*, *CD83* and *REL* (Extended Data Fig. 6e), which prompted us to investigate the presence of follicular dendritic cells (FDC). Indeed, we detected CD21^+^ FDC in advanced and tumour-stage CTCL samples using IHC (Extended data Fig. 6f,g). In addition, we detected the expression of genes associated with follicular helper T cells and B cell recruitment (i.e., *BCL6, PDCD1* and *CXCL13*) expressed by malignant T cells (Extended data Fig. 6h). CTCL fibroblasts also share expression features with TRC in healthy lymph nodes^39^, which may indicate a role in TLS formation.

Spatial mapping and multi-colour immunofluorescence imaging showed proximity and direct cell-cell contact between CD20^+^ B cells and CD4^+^ tumour cells in CTCL TME (Fig. 4d,e, Extended data Fig. 6i,j). We next performed cell-cell interaction analysis and identified putative ligand-receptor interactions between B cells and tumour cells, which included costimulatory interactions such as CD70-CD27, CD40LG-CD40, CD58-CD2 and ALCAM-CD6 that are known to promote T cell activation, and B cell recruitment interaction CXCL13-CXCR5 that is related to lymphoid structure formation (Fig. 4f).

As these observations together suggest a role of B cells in promoting tumour growth, we investigated the correlation of B cell abundance in CTCL lesional skin to patient clinical outcome data. The abundance of B cells inferred from bulk RNA-seq data tended to associate with poor disease prognosis (Fig. 4g, Extended data Fig. 6k). In accordance with this, we detected increased percentages of B cells in progressive CTCL skin lesions using IHC (Fig 4h, Extended data Fig. 6l). Interestingly, three patients from the Vienna cohort, who died from CTCL, displayed increased B cell presence in tumour samples (IIB stage at biopsy) taken 3 to 7 years before death (Fig. 4h, Extended data Fig. 6l). These findings support the utility of B cell immunostaining as a potential diagnostic and prognostic aid for CTCL particularly in early-stage disease.

We further performed drug2cell^46^ analysis, which predicts therapeutic targets leveraging the ChEMBL database, and identified several known CD20-directed antibodies (rituximab, obinutuzumab) as well as elotuzumab (approved for treatment of multiple myeloma) to target CTCL-associated B cells and malignant T cells (Fig. 4i, Supplementary Table 7). These data provide evidence for B cells as a therapeutic target in CTCL. Indeed, there have been isolated reports of CTCL patients responding to incidental or intentional treatment with rituximab^47,48^. In addition, drug2cell analysis also identified 15-hydroxyprostaglandin dehydrogenase (HPGD) as a potential selective drug target against malignant T cells and sparing benign T cells.

Taken together, our data revealed a trajectory of malignant T cells to co-opt Th2-like gene programmes and supported by a Th2 permissive pro-tumorigenic TME in CTCL moving from early stage to advanced stage disease. Notably, we demonstrate B cells forming aggregates and tertiary lymphoid-like structures, which are associated with disease progression and outcome (Fig. 4g,h,j, Extended data Fig. 6m).

## Discussion

CTCL exhibits a wide spectrum of genetic and clinical alterations with limited specific histological features in early stages, impeding diagnosis. Our findings reveal metabolically altered clonal Th2-like malignant cells in a Th2-permissive tumour-promoting microenvironment mainly contributed by MHC-II^+^ fibroblasts and monocyte-derived dendritic cells. Th2-like malignant T cells are also associated with B cell infiltration and aggregate formation which can be used to aid CTCL diagnosis and potentially treatment.

Our in-depth characterisation of malignant T cell clones revealed extensive CNVs across their genomes. Despite high inter-patient heterogeneity, which had also been reported in previous CTCL transcriptome studies^9,16,21^, we found metabolic features that were conserved across tumours in our cohort and observed elsewhere in cells in hypoxic environments (the Warburg effect) as well as circulating memory T cells^34^. Glycolytic metabolism is associated with distinct cellular architecture, including mitochondrial polarisation that results in altered mitochondrial structure. Interestingly, such altered structures have been observed in electron microscopic analysis of skin T cells in MF^49^. In addition to CTCL-specific meta-programmes, we identified molecules that were not ubiquitously expressed by all tumours, but could be developed for diagnostic use collectively, such as by multiplexed targeted transcriptome profiling. This includes the DNA-binding protein TOX, which has been proposed as a potential but not exclusive marker for CTCL^50^, and GTSF1, a protein whose function in T cells is unknown and is identified in some CTCL cancer cell lines^51^.

Surprisingly, differential abundance testing revealed a predominance of B cells in CTCL compared to healthy skin, AD and psoriasis, which was corroborated by bulk deconvolution, spatial transcriptional profiling, immunofluorescence and IHC staining. B cells rarely appear in healthy skin, AD and psoriasis, and thus their presence in CTCL is significant. Importantly, our analyses reveal that B cell abundance is associated with progressive disease and poor prognosis, in keeping with previous case reports^52^. Dissecting the role of B cells in CTCL revealed interactions with malignant T cells, as well as the presence of GC-like B cells and the formation of B cell and follicular dendritic cell aggregates resembling tertiary lymphoid structures. This is supported by previous dermato-pathological studies which identified the expression of follicular helper T cell markers in CTCL^53^. Importantly, B cell infiltration in classical CTCL entities needs to be discerned from rare subtypes of cutaneous non-Hodgkin lymphomas, characterised by aberrant expression of CD20 in malignant T cells^54^. Collectively, our data suggests that B cell-depleting therapies may effectively target the CTCL TME, providing evidence for wider use of these therapies in CTCL. The interaction we identified between malignant T cells and MHC-II^+^ fibroblasts via the CCR4-CCL5 axis may also explain the therapeutic efficacy of mogamulizumab in CTCL^17^.

A Th2-TME has been found to foster tumour growth in non-hematopoietic malignancies including breast and pancreatic cancer^55,56^. Here we show that malignant T cells in CTCL co-opt a Th2-immune programme to promote recruitment of B cells and tertiary lymphoid structures in the skin, a non-lymphoid tissue. The Th2 microenvironment may in turn promote the survival of malignant T cells. Whether the Th2-immune programme deployment is aided by an antigen specific (including response to a skin microbe) or antigen non-specific manner remains to be determined. Interestingly, the blocking antibody to IL-4 receptor, dupilumab, has been shown to unmask (CTCL misdiagnosed as AD) or worsen CTCL symptoms^57^, likely by increasing free IL-4 and IL-13 to bind to IL-13α2 receptor^58,59^.

In summary, our findings provide a new understanding of CTCL malignant cells within their TME, including the co-optation of Th2-immune programme resulting in B cell aggregates and tertiary lymphoid structure formation in advanced stage disease. These findings provide evidence to support the deployment of B-cell directed (combination-)therapies to treat patients with CTCL.

---

**Extended Data Fig. 1.**
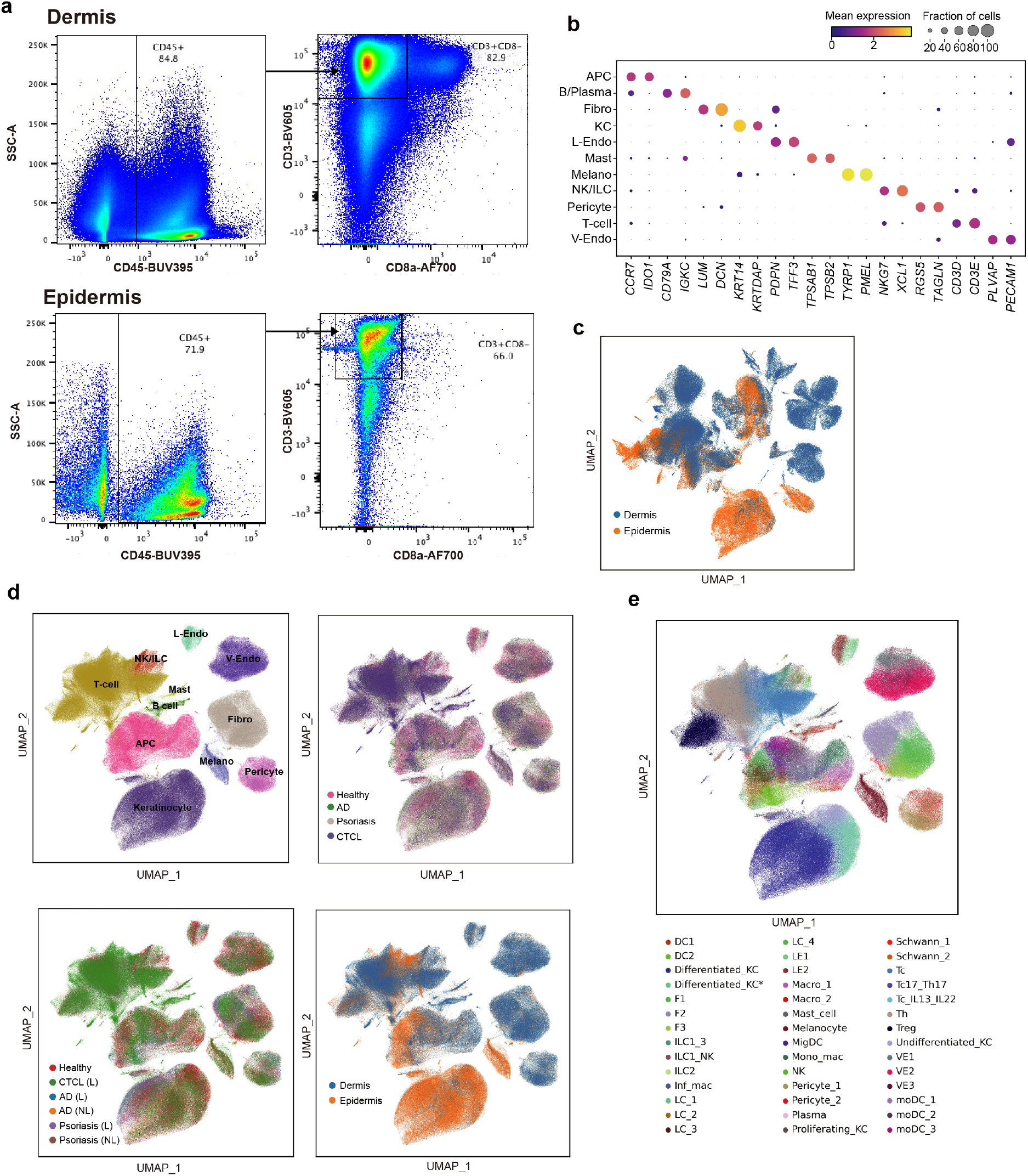
**a,** Representative FACS gating strategy for scRNA-seq for dermal (top) and epidermal (bottom) samples. Plots follow on from classical live singlet gating using DAPI and FSC-A/FSC-H/SSC-W respectively. **b**, Gene expression dot plot of marker genes for broad cell types. Dot colour indicates log-transformed and normalised expression value. Dot size indicates the percentage of cells in each cell type expressing a given gene. **c**, Overall UMAP of our CTCL dataset with cells coloured by the skin layers they derive from. **d**, UMAPs showing the integration of skin cell atlas data, coloured by broad cell types, skin conditions, lesion or non-lesion, and skin layers. **e**, UMAP showing cell type predictions through logistic regression-based label transfer based on skin cell atlas.

**Extended Data Fig. 2.**
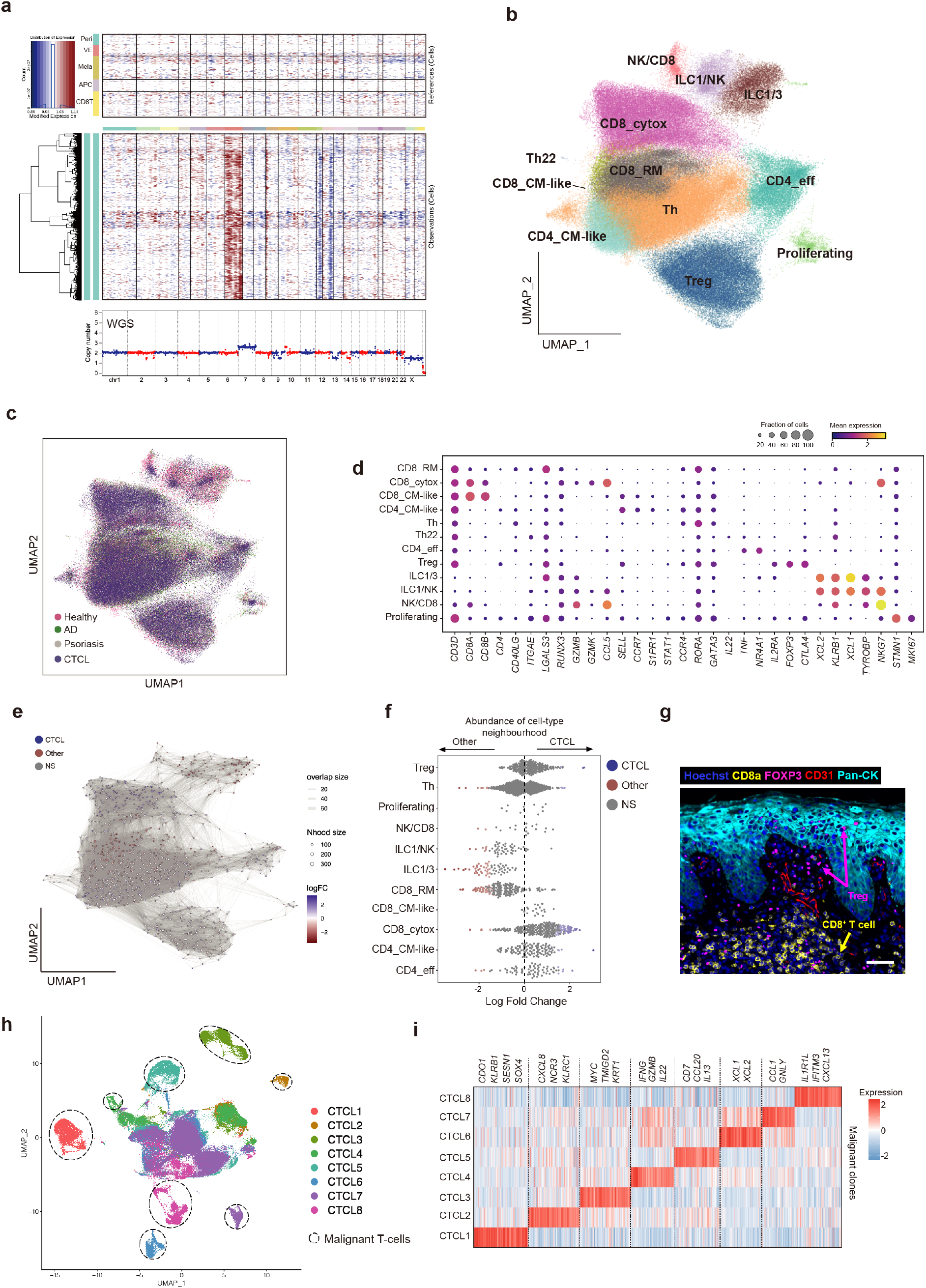
**a**, Upper: Heat map showing CNVs inferred from scRNA-seq data in a representative sample (Donor 1). Top panel represents reference normal cells. Peri, pericytes; VE, vascular endothelial cells; Mela, melanocytes. Bottom heat map represents malignant T cells. Lower: CNVs inferred from whole genome sequencing of the same sample. **b** and **c**, UMAP visualisation of the benign T cell and NK/ILC population in CTCL dataset integrated with skin cell atlas, coloured by cell types in **b** and skin conditions in **c**. **d**, Gene expression dot plot of marker genes for T cell and NK/ILC subsets. **e**, Neighbourhood graph of the results from Milo differential abundance testing. Nodes are neighbourhoods, coloured by their log fold change across ages. Non-differential abundance neighbourhoods (FDR 10%) are coloured white, and sizes correspond to the number of cells in each neighbourhood. Edges depict the number of cells shared between neighbourhoods. **f**, Beeswarm plot of the log-transformed fold changes in abundance of cells in CTCL versus those in healthy skin, AD and psoriasis (as Other in the plot) from skin cell atlas. Differential abundance neighbourhoods at FDR 10% are coloured. NS, not significant. **g**, Multi-colour immunofluorescence image (Rarecyte) in a representative tumour. Scale bar, 50 μm. **h**, UMAP of T, NK and ILC cells in our CTCL dataset coloured by donors. Dashed circles highlight malignant T cell clones. **i**, Heat map showing inter-tumour DEGs across the eight donors. Representative genes are labelled.

**Extended Data Fig. 3.**
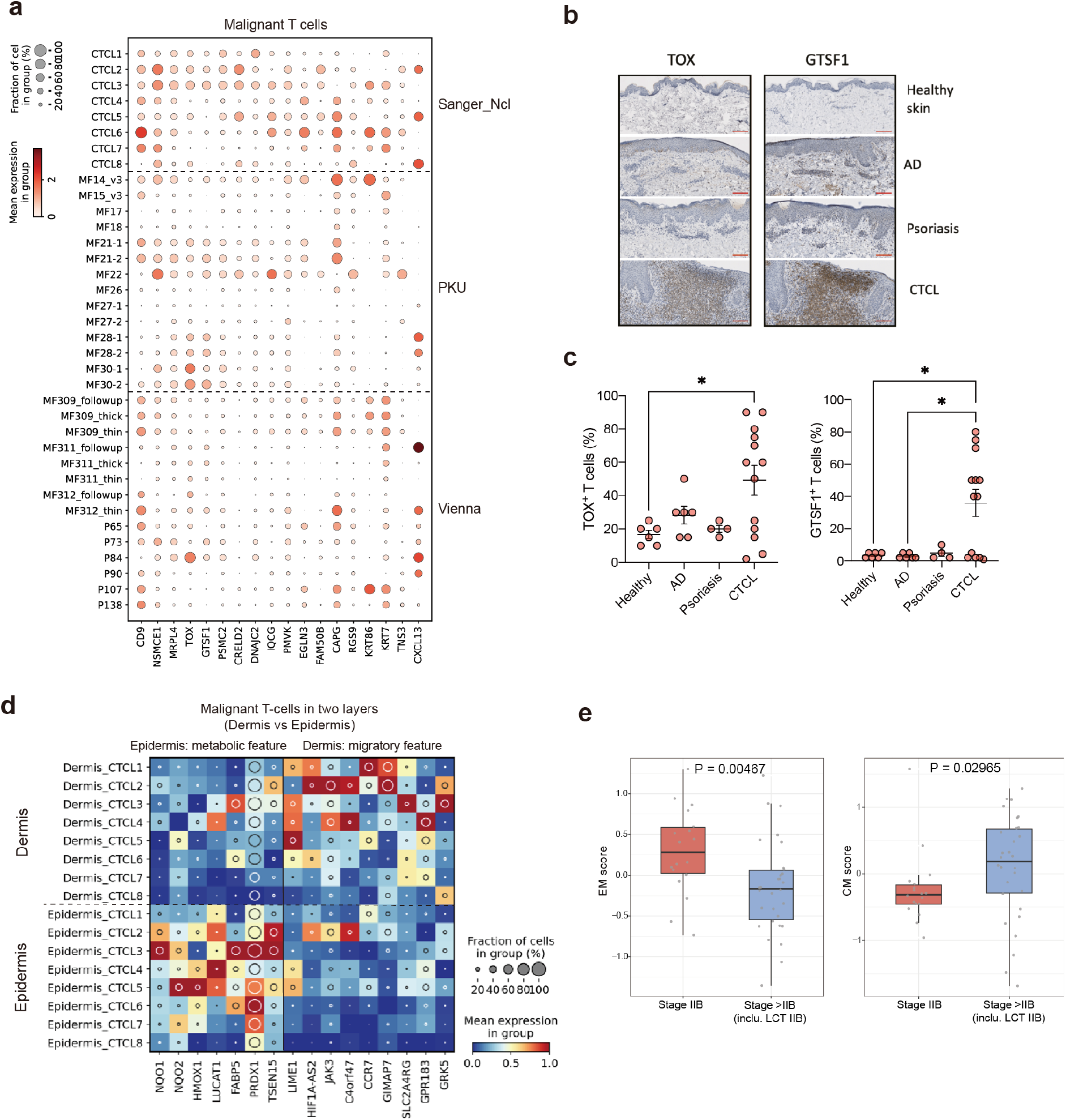
**a**, Gene expression dot plot of genes upregulated in malignant T cells in our dataset and other two published datasets (Vienna^16^ and PKU^17^). **b**, IHC staining for TOX and GTSF1 in representative healthy skin, AD, psoriasis and CTCL samples. Scale bar, 100 µm. **c**, Dot plots comparing the IHC results in healthy skin, AD, psoriasis and CTCL samples. The asterisk represents p<0.05, one-way anova and Tukey’s multiple comparison test **d**, Heat map of DEGs between malignant T cells from dermis and epidermis. Colour represents expression level standardised between 0 and 1. The inset circle indicates the percentage of cells expressing a given gene. **e**, Box plots showing the expression of two gene scores in bulk RNA-seq data. Patients are divided into two groups: 1. stage IIB (without large cell transformation: LCT; n=18) and over stage IIB plus LCT (n=30). EM, effector memory; CM, central memory. Gene lists for calculating scores are shown in Fig. 2e. The lower edge, upper edge and centre of the box represent the 25th (Q1) percentile, 75th (Q3) percentile and the median, respectively. The interquartile range (IQR) is Q3 – Q1. Outliers are values beyond the whiskers (upper, Q3 + 1.5 × IQR; lower, Q1 − 1.5 × IQR).

**Extended Data Fig. 4.**
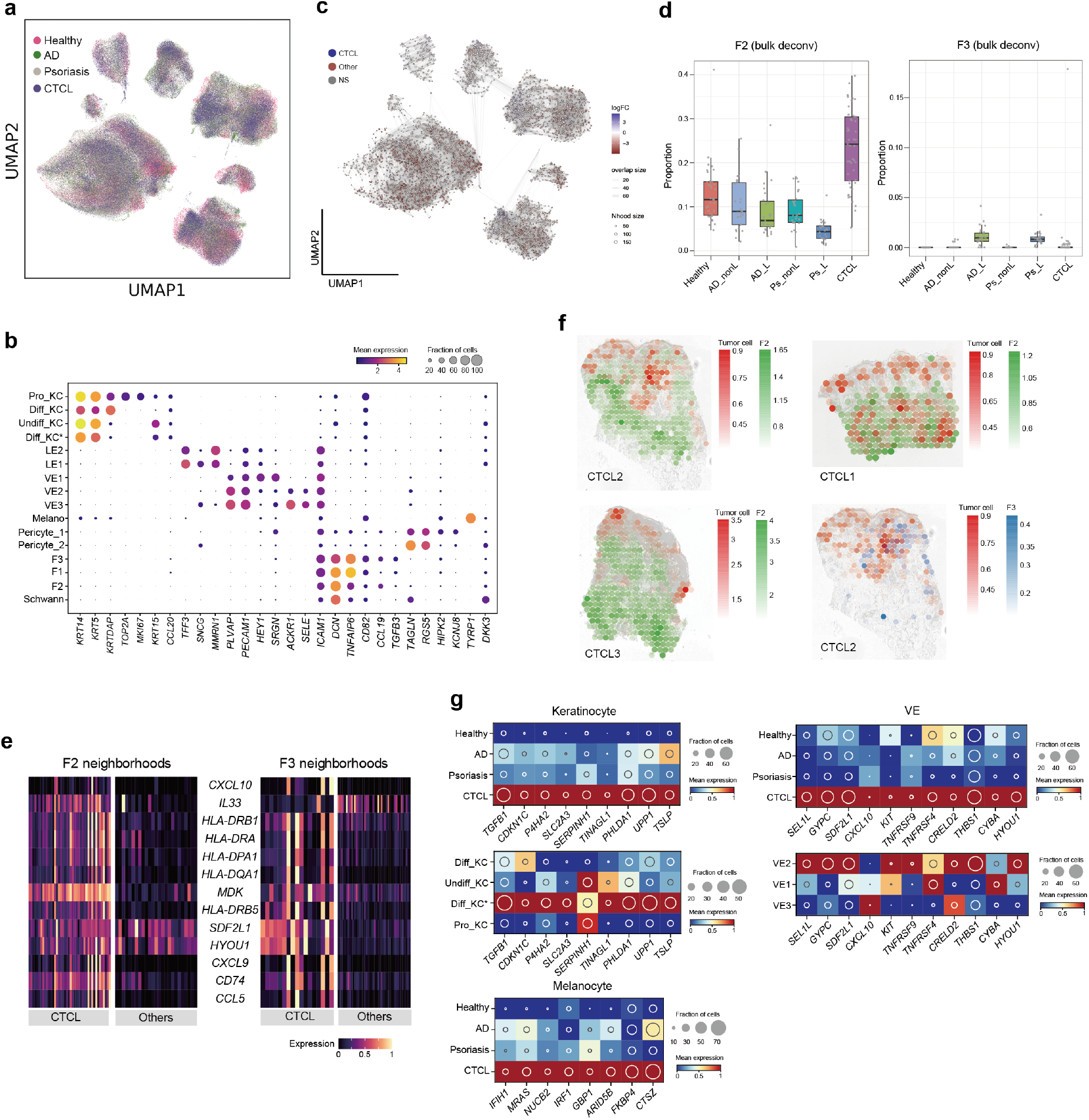
**a**, UMAP visualisation of the stromal population in the current dataset integrated with skin cell atlas, coloured by skin conditions. **b**, Gene expression dot plot of marker genes for stromal cell subsets. Dot colour indicates log-transformed and normalised expression value. Dot size indicates the percentage of cells in each cell type expressing a given gene. **c**, Neighbourhood graph of the results from Milo differential abundance testing. Nodes are neighbourhoods, coloured by their log fold change across ages. Non-differential abundance neighbourhoods (FDR 10%) are coloured white, and sizes correspond to the number of cells in each neighbourhood. Edges depict the number of cells shared between neighbourhoods. **d**, Box plot of deconvolution of F2 and F3 in bulk RNA-seq datasets of healthy skin (n=38), AD (nonlesion: n =27; lesion: n=27), psoriasis (nonlesion: n =27; lesion: n=28) and CTCL (n=49). Numbers of samples in the categories are labelled. The lower edge, upper edge and centre of the box represent the 25th (Q1) percentile, 75th (Q3) percentile and the median, respectively. The interquartile range (IQR) is Q3 – Q1. Outliers are values beyond the whiskers (upper, Q3 + 1.5 × IQR; lower, Q1 − 1.5 × IQR). **e**, Heat map showing the expression of genes in Fig. 3c in CTCL enriched cell neighbourhoods compared to other three conditions. **f**, Spatial mapping of F2 and tumour cells in three representative samples and F3 and tumour cells in one representative sample. Estimated abundance (colour intensity) is overlaid on histology images. **g**, Heat map of DEGs in CTCL enriched keratinocyte, vascular endothelial cell and melanocyte. Colour represents expression level standardised between 0 and 1. The inset circle indicates the percentage of cells expressing a given gene.

**Extended Data Fig. 5.**
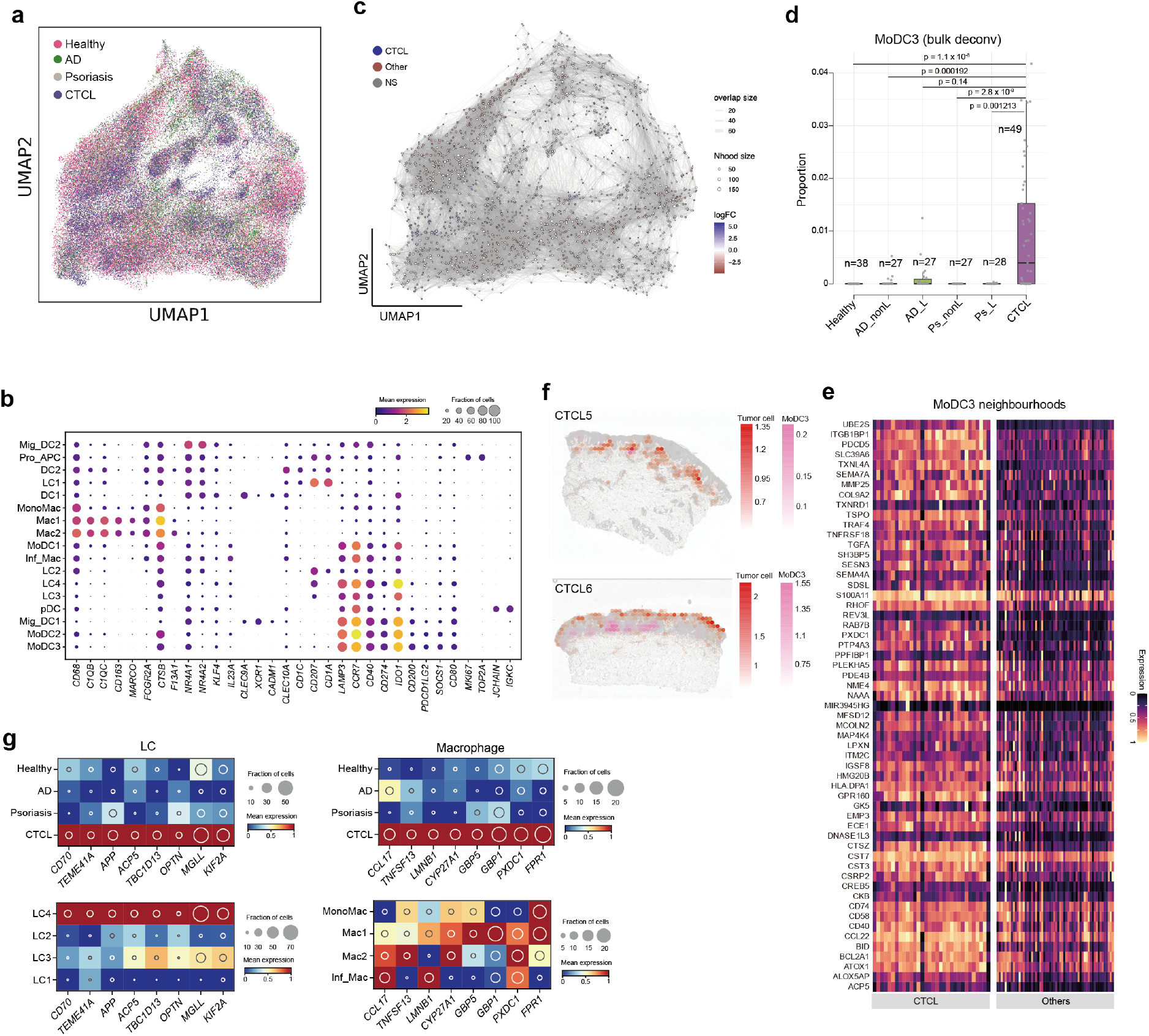
**a**, UMAP visualisation of the APC population in the current dataset integrated with skin cell atlas, coloured by skin conditions. **b**, Gene expression dot plot of marker genes for APC cell subsets. Dot colour indicates log-transformed and normalised expression value. Dot size indicates the percentage of cells in each cell type expressing a given gene. **c**, Neighbourhood graph of the results from Milo differential abundance testing. Nodes are neighbourhoods, coloured by their log fold change across ages. Non-differential abundance neighbourhoods (FDR 10%) are coloured white, and sizes correspond to the number of cells in each neighbourhood. Edges depict the number of cells shared between neighbourhoods. **d**, Box plot of deconvolution of MoDC3 in bulk RNA-seq datasets of healthy skin (n=38), AD (nonlesion: n =27; lesion: n=27), psoriasis (nonlesion: n =27; lesion: n=28) and CTCL (n=49). Two-sided Wilcoxon rank-sum. The lower edge, upper edge and centre of the box represent the 25th (Q1) percentile, 75th (Q3) percentile and the median, respectively. The interquartile range (IQR) is Q3 – Q1. Outliers are values beyond the whiskers (upper, Q3 + 1.5 × IQR; lower, Q1 − 1.5 × IQR). **e**, Heat map showing DEGs in CTCL enriched MoDC3 neighbourhoods. Other, healthy/AD/psoriasis. **f**, Spatial mapping of MoDC3 and tumour cells in three representative samples. Estimated abundance (colour intensity) is overlaid on histology images. **g**, Heat map of DEGs in CTCL enriched Langerhans cell and macrophage. Colour represents expression level standardised between 0 and 1. The inset circle indicates the percentage of cells expressing a given gene.

**Extended Data Fig. 6.**
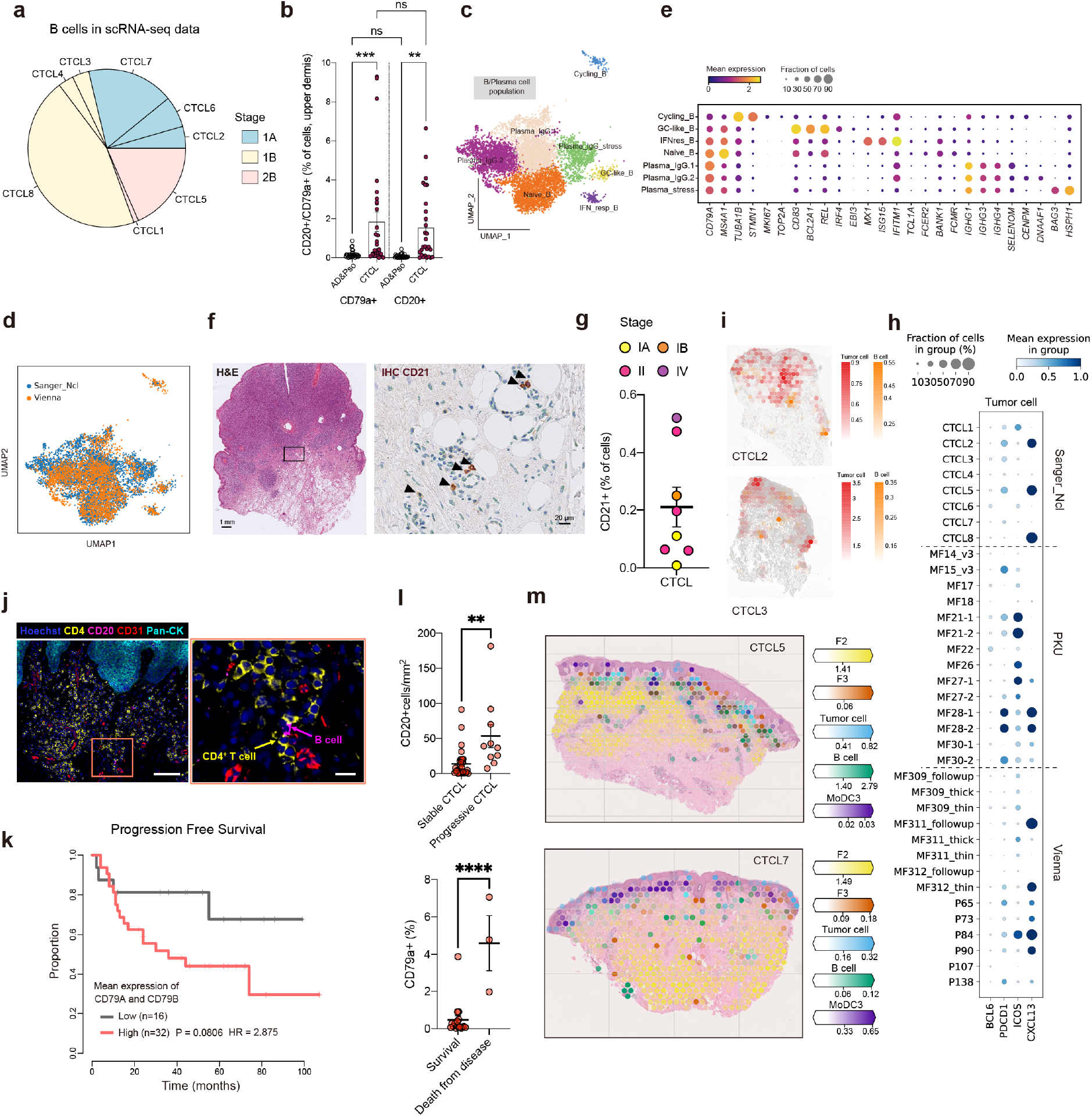
**a**, Pie chart showing proportions of B cells from each of the eight donors in our CTCL dataset. Colours indicate different disease stages. **b**, CD79a^+^ and CD20^+^ cells in CTCL (n=27, Newcastle cohort) and AD/Pso (n=30). Data shown as individual values and mean percentage of IHC+ cells among all cells of the dermis, error bars indicate SEM. **c** and **d**, UMAPs of B cell population in integrated data (integrated with Vienna dataset^16^) coloured by cell subsets in **c** and datasets in **d**. **e**, Gene expression dot plot of marker genes for B cell subsets. Dot colour indicates log-transformed and normalised expression value. Dot size indicates the percentage of cells in each cell type expressing a given gene. **f**, Representative H&E staining (left) and CD21 IHC (right) of an advanced tumour-stage CTCL sample. Arrows show CD21^+^ FDC. Scale bars, 1 mm and 20 μm. **h,** Dot plot showing the expression of follicular T cell marker genes in malignant T cells in three scRNA-seq datasets (Sanger_Ncl: current dataset, PKU^17^, and Vienna^16^). **g**, Percentages of CD21^+^ cells in CTCL with different stages. **i**, Spatial mapping of B cell and tumour cells in two representative samples. Estimated abundance (colour intensity) is overlaid on histology images. **j**, Multi-colour immunofluorescence images (Rarecyte) in a representative tumour. A representative view showing B cell and CD4^+^ T cell interaction is zoomed in. Scale bars, 100 μm and 20 μm. **k**, Progression free survival probability of CTCL patients according to stratification of B cell abundance estimated by mean expression of *CD79A* and *CD79B*. HR, hazard ratio. **l**, IHC staining of CD20 in stable and progressive CTCL skin samples (upper) and the staining of CD79a and association with outcome (survival vs. death from disease, lower). Data shown as individual values and mean percentages of CD20^+^ cells per mm^2^ +/- SEM (upper) and individual values and mean percentages of CD79a^+^ cells among all cells +/- SEM, n=27 (Vienna cohort), **, P<0.01, ****, P<0.0001. **m**, Spatial mapping of tumour cells, F2, F3, MoDC3 and B cell in Visium data for two representative tumours. Estimated abundance for cell types (colour intensity) across locations (dots) is overlaid on histology images using cell2location.

## Online Methods

### Patient recruitment and sample acquisition

Skin samples generated for this study, from patients with CTCL were donated with written consent and approval from the Newcastle and North Tyneside NHS Health Authority Joint Ethics Committee (08/H0906/95+5). Each CTCL patient donated two skin punch biopsies, from a representative plaque or tumour. One biopsy was used for scRNA-seq, and the other biopsy for bulk sequencing and IHC. All patients had MF, diagnosed based on correlation of clinical and histopathological features. Stage of CTCL at time of biopsy was taken from the patients notes and based on clinical assessment performed by dermatology specialists at the Department of Dermatology and NIHR Newcastle Biomedical Research Centre, Newcastle, UK, using the Modified Severity-Weighted Assessment Tool (mSWAT). For additional IHC validation cohorts, samples were donated with consent from the local ethics committee at the Medical University of Vienna (ECS 1360/2018) and the Swedish Ethical Review Authority (2019-03467). CTCL diagnosis and staging as well as monitoring for disease progression was performed by specialists in dermatology and dermato-histopathology at the Department of Dermatology, Medical University of Vienna and the Department of Dermatology, Karolinska University Hospital, Stockholm.

### Sample processing

Skin biopsies were immediately processed by removing the lower dermis and subcutis and separating epidermis and dermis after dispase II digestion at a concentration of 2U/ml for 2-3 hours at 37°C. Epidermis and dermis were processed separately in type IV collagenase at a concentration of 1.6 mg/ml overnight (37°C 5% CO2). Subsequently, single cell suspensions were formed by vigorous pipetting and filtering (100-micron filter), counted and further processed via FACS.

### FACS sorting and 10x Genomics Chromium loading

Both cells from the epidermis and dermis were stained with an antibody panel containing CD45 (BD Biosciences) and CD8a (Biolegend) and sorted using FACS into the following fractions: CD45^-^, CD45^+^ CD8a^+^ and CD45^+^ CD8a^-^. A target of 10,000 cells was used to calculate the loading volume for the 10x Chromium, taken from the manufacturer’s protocol. Each fraction sorted from the epidermis and dermis was loaded onto one channel of the 10x Chromium chip before running on the Chromium Controller using the 10x 5’ v1 kits.

### Library preparation and sequencing

Gene expression libraries were generated from the resulting cDNA after clean up following the 10x Genomics protocols. Enriched TCR cDNA was also generated from each CD45+ fraction and subsequent libraries were made. All libraries were sequenced using an Illumina NovaSeq with the gene expression libraries sequenced to achieve a minimum of 50,000 reads per cell and the TCR libraries sequenced to achieve a minimum of 5,000 reads per cell.

### Whole genome sequencing

A small piece of each skin biopsy was frozen at -20℃ in RNA Later (Invitrogen) for 24 hours before the liquid was removed and the samples moved to -80℃. DNA was extracted from frozen skin samples using the AllPrep Micro kit (Qiagen) following the manufacturer’s protocol. The DNA was quantified using a Qubit with the High Sensitivity DNA kit (Invitrogen). Library preparation was carried out using NEBNext® Ultra™ II DNA Library Prep Kit from Illumina. Libraries were uniquely dual indexed to mitigate for tag hopping. Libraries were then quantified and equimolar pooled. The pool was sequenced down 1 lane of an S4 flow-cell on the Illumina NovaSeq 6000 platform, with 150bp paired end reads.

### FFPE Visium CytAssist spatial transcriptomics

RNA quality and tissue morphology of the CTCL formalin-fixed and paraffin embedded (FFPE) sample blocks were assessed prior to FFPE Visium processing. Each of the eight CTCL FFPE sample blocks was sectioned using a microtome (Leica RM2235) at 5 um thickness onto a SuperFrost Plus microscope slide (VWR, 6310108), incubated for 3 hours at 42℃, dried overnight in a dessicator at room temperature and processed for FFPE Visium within 2 weeks of sectioning. Deparaffinization, hematoxylin and eosin (H&E) staining and decrosslinking steps were performed as per manufacturer’s recommendations (10x Genomics Demonstrated Protocol, CG000520) and sections were imaged on a Hamamatsu Nanozoomer. Sections were then further processed with FFPE Visium CytAssist v2 chemistry (6.5 mm) kit and dual-indexed libraries were prepared as per 10x Genomics User Guide, CG000495. Four libraries were pooled at a time and sequenced down one lane of Illumina Novaseq SP flow cell with the following run parameters: read 1: 28 cycles; i7 index: 10 cycles; i5 index: 10 cycles; read 2S: 50 cycles.

### RareCyte 16-plex immunofluorescence staining

All steps were performed at room temperature unless stated otherwise. Briefly, FFPE sample blocks were sectioned using a microtome (Leica RM2235) at 5 µm thickness and placed on a superfrost slide (Fisher scientific 12312148). Slides were dried at 60°C for 60 min to ensure tissue sections had adhered to the slides. After deparaffinization, tissue sections were subjected to antigen retrieval using the BioGenex EZ-Retriever system (95°C for 5 min followed by 107°C 5 min). To remove autofluorescence, slides were bleached with AF Quench Buffer which consists of 4.5% H2O2 / 24 mM NaOH in PBS. Slides were quenched for 60 min using the HIGH setting with a strong white light exposure followed by further quenching for 30 min using 365 nm HIGH setting using a UV transilluminator. Slides were rinsed with 1X PBS and incubated in 300 µl of Image-iT™ FX Signal Enhancer (Thermo Fisher, # I36933) for 15 min. Slides were rinsed and 300 µl of labelled primary antibody staining cocktail was added to the tissue, which subsequently was incubated for 120 min in the dark within a humidity tray. All antibodies were pre-diluted according to company recommendations and were not adjusted further. Details about antibodies used can be found in Supplementary table 8. Slides were washed with a surfactant wash buffer and 300 µl of nuclear staining in goat diluent was added to the slide. Slides were then incubated in the dark for 30 min in a humidity tray. Slides were then washed and placed in 1X PBS. Finally, the slides were coverslipped using ArgoFluor mount media and left in the dark at room temperature overnight to dry. Slides were imaged on the following day using a RareCyte Orion microscope with a 20X objective. Scans were performed using Imager and relevant acquisition settings were applied using the software Artemis. Slides were subsequently transferred to -20°C for extended storage.

### Single cell RNA-seq data processing, quality control and doublet removal

Gene expression and VDJ data from droplet-based sequencing were processed using the 10x software package CellRanger (version 3.1.0 and vdj) and aligned to the GRCh38 reference genome (official Cell Ranger reference, version 3.0.0). Gene expression outputs from CellRanger were read in using the read_10x_mtx function in Scanpy^60^ (version 1.8.1). Data objects from different 10x lanes were then concatenated using the concatenate function in anndata (version 0.7.6). To detect and remove doublets, we applied Scrublet^61^ (version 0.2.3) to the data from each 10x lane to obtain per-cell scrublet scores and used a doublet exclusion threshold of median plus four median absolute deviations of the doublet score, as previously described^22^. Cells with greater than 20% mitochondrial gene expression or expression of fewer than 200 detected genes were excluded from downstream analysis. Genes that were expressed in fewer than 3 cells were also removed.

### Data normalisation, embedding, visualisation, clustering and integration

We further performed data normalisation to correct for cell-to-cell variation using the normalize_per_cell function in Scanpy (version 1.8.1). Normalised data were then transformed using the log1p function in Scanpy to alleviate skewness of data and mean-variance relationship. Expression values of each gene were then scaled and centred using the scale function in scanpy. Highly variable genes (HVGs) were detected using the highly_variable_genes function in scanpy with minimum cut-off values 0.0125 and 0.5 for expression and dispersion. We then regressed out percentages of mitochondrial gene expression using the regress_out function and further calculated principal components (PCs) using the pca function in scanpy. For batch correction, we used the harmonypy^62^ package (https://github.com/slowkow/harmonypy) to correct donor-to-donor variation with the theta value being set to 3. The neighbours function was used to calculate the neighbourhood graph. UMAP embedding was calculated using the umap function in scanpy. The neighbourhood graph was then clustered using the leiden function in scanpy. Broad cell types were annotated based on expression of canonical marker genes. We integrated our CTCL dataset with the skin cell atlas dataset and two published CTCL datasets. First, datasets were concatenated using the concatenate function in anndata. The downstream data normalization process was the same as mentioned above. We ran harmonypy for batch correction using donor as the batch key and setting theta value to 3.

### Differential abundance analysis using Milo

To reveal potential differences in cellular abundance in CTCL, we performed differential abundance analysis comparing CTCL to healthy skin, AD and psoriasis using Milo^26^. For the overall integrated object and different major cell compartments, we first performed a random subsampling using the subsample function in Scanpy, which subsampled the overall object to 0.1, stromal population to 0.3, APC population to 0.5, and benign T-cell population to 0.3 of the total numbers of cells. Then the standard Milo pipeline was run for each data object with the proportion of graph vertices to randomly sample (prop in the makeNhoods function) being set to 0.05, k being set to 20, and d being set to 30. Beeswarm plots were made to show the log-transformed fold changes in abundance of cells in CTCL versus those in healthy skin, AD and psoriasis for each data object.

### Inferring copy number variations based on scRNA-seq data

To effectively distinguish malignant T cells and non-malignant cells, we inferred large-scale chromosomal copy number variations of single cells based on scRNA-seq data using the tool InferCNV (https://github.com/broadinstitute/inferCNV) with default parameters. Briefly, InferCNV first orders genes according to their genomic positions (first from chromosome 1 to X and then by gene start position) and then uses a previously described sliding-average strategy to normalise gene expression levels in genomic windows with a fixed length. Multiple putative non-malignant cells are chosen as the reference to further denoise the CNV result.

### Analysing intra-tumour expression programmes and meta-programmes

In order to explore intra-tumour expression programmes, we applied non-negative factorization (implemented in the R NMF package) to the tumour cells from the eight CTCL patients. Briefly, for each tumour, we first normalised the expression counts using the NormalizeData function in Seurat with default parameter settings. Highly variable genes (HVGs) were then selected using the FindVariableFeatures function in Seurat^63^. Next, we performed centre-scale for HVSs and regressed out the percentage of mitochondria genes using the ScaleData function. For NMF analysis, all negative values in the expression matrix were replaced by zero. The top 10 ranked NMF gene modules in each tumour sample were extracted using the nmf function in the NMF package. For each gene module, we extracted the top 30 genes with the highest weight which were used to define a specific intra-tumour expression programme. Finally, we only included intra-tumour expression programmes that had standard deviations larger than 0.1 among tumours cells. To investigate if some intra-tumour expression programmes were actually shared by multiple tumours, we applied a clustering analysis to all programmes based on the pair-wised Jaccard index calculated as follows, where A and B represent two intra-tumour programmes.

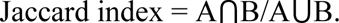

We defined those intra-tumour programmes shared by multiple tumours as meta-programmes (MPs). For those MPs that consist of more than two intra-tumour programmes, we used genes shared by at least 50% intra-tumour programmes to define them. While for those MPs that consist of two intra-tumour programmes, we used genes shared by the two intra-tumour programmes to define them.

### Sub-clustering and annotation of different cell compartments

We performed sub-clustering and annotation of different cell compartments based on objects integrated with the skin cell atlas data. To facilitate the annotation, we first trained a logistic regression (LR) model using the skin cell atlas data as training data and predicted identities of cells in our CTCL dataset. Cells were then subset into stromal, APC, and benign T/NK/ILC population based on the expression of cell lineage markers and clustering results. We then performed data normalisation, embedding, visualisation and clustering on each cell population. For each cell population, we regressed out mitochondrial gene percentage using the regress_out function in Scanpy. For the APC population, we additionally regressed out ribosomal gene percentage. Donor-to-donor variation was corrected using the harmonypy package with the theta value being set to 3. For each cell population, we performed clustering using the leiden function in Scanpy and manually annotated clusters based on LR predicted cell identities and differentially expressed genes.

### Visium data processing and spatial mapping of cell types with cell2location

Sequencing reads from 10x Genomics Visium FFPE libraries were aligned to the human transcriptome reference GRCh38-2020-A using 10x Genomics SpaceRanger (v.2.1.0) and exonic reads were used to produce mRNA count matrices for each sample. 10x Genomics SpaceRanger was also used to align paired histology images with mRNA capture spot positions in the Visium slide. To spatially map the cell types annotated in scRNA-seq data to their spatial locations in tissues, we applied cell2location to integrating scRNA-seq data of CTCL with Visium FFPE mRNA count matrices as described previously^27^. Briefly, the cell2location model estimates the abundance of each cell type in each location by decomposing mRNA counts in Visium FFPE data using the transcriptional signatures of reference cell types. Two major steps were in analysis using cell2location: (1) We applied a negative binomial regression model implemented in cell2location and estimated the reference signature of fine-grained annotated cell types in scRNA-seq data. In this step, we used an unnormalized mRNA count matrix as input and filtered it to 13,581 genes and 279,561 cells. Donor IDs were regarded as the batch category and the following parameters were used to train the model: ‘max_epochs’ = 500, ‘batch_size’ = 2500, ‘train_ size’ = 1 and ‘Ir’ = 0.002. (2) The reference signature model was further used by cell2location to estimate spatial abundance of cell types. We kept genes that were shared with scRNA-seq and estimated the abundance of cell types in the eight Visium FFPE samples. In this step, cell2location was used with the following parameter settings: training iterations: 20,000, number of cells per location N = 7, ‘detection_alpha’ = 20. We to estimate cell type abundance in dependence on distance to surface, we first normalised cell type abundance by dividing it by per-spot totals. Then we grouped spots by rounded distance to surface and calculated mean and standard deviation of mean for each cell type and each distance.

### Differentially expressed gene analysis using a pseudo-bulk strategy

We applied a pseudo-bulk strategy to the analysis of differentially expressed genes (DEGs) between (1) malignant and benign T cells, (2) malignant T cells from epidermis and dermis, (3) malignant T cells from early stage and advanced stage samples, and (4) microenvironmental cells from CTCL and other three conditions. Briefly, we aggregated raw counts of each gene by donor and used donors rather than cells as biological replicates. DEG analyses were carried out using R package edgeR^64^. For the analysis of (1), we excluded non-lesion cells from AD and psoriasis and regarded healthy skin, AD and psoriasis as one comparator (other). We filtered genes by expression levels using the filterByExpr function in edgeR with ‘min.count’ and ‘min.total.count’ being set to 50 and 100 respectively. We designed the model matrix using the model.matrix function and only included one variable, namely groups (malignant T cell and benign T cell). For the analysis of (2), we first divided malignant T cells from each patient into those from epidermis and dermis, and conducted pseudobulk on tissue plus patient (i.e., CTCL1_dermis and CTCL1_epidermis were aggregated separately). In model matrix design, we fit the model on paired samples considering both tissue (dermis and epidermis) and patient (CTCL1 to CTCL8). For the analysis of (3), we included both studies (PKU, Vienna and Ncl_Sanger) and groups (malignant T cell and benign T cell) as variables to consider variation across studies. For the analysis of (4), we excluded non-lesion cells from AD and psoriasis and regarded healthy skin, AD and psoriasis as one comparator (other). We filtered genes by expression levels using the filterByExpr function in edgeR with ‘min.count’ and ‘min.total.count’ being set to 50 and 100 respectively. We designed the model matrix using the model.matrix function and only included one variable, namely groups CTCL and other (healthy skin, AD and Psoriasis). For all the analysis, we fit genewise negative binomial generalised linear Models with quasi-likelihood tests using the glmQLFit and glmQLFTest functions in edgeR.

### Bulk deconvolution of cell types in healthy skin, AD, psoriasis and CTCL

For bulk deconvolution analysis, we first downloaded published bulk RNA-seq datasets of healthy skin, AD, psoriasis, and CTCL from the Gene Expression Omnibus (GEO) database with the accession codes GSE121212 and GSE168508. A single-cell reference for deconvolution analysis was then prepared by randomly downsampling the integrated object (healthy skin, AD psoriasis, and CTCL) to 8% of total cells. BayesPrism^36^ was used for deconvolution analysis with raw counts for both single-cell and bulk RNA-seq data as inputs. Both the ‘cell type labels’ and the ‘cell state labels’ were set to fine-grained annotations. Ribosomal protein genes and mitochondrial genes were removed from single-cell data as they are not informative in distinguishing cell types and can be a source of large spurious variance. We also excluded genes from sex chromosomes and lowly transcribed as recommended by the BayesPrism tutorial. For further analysis, we applied a pairwise t-test to select differentially expressed genes with the ‘pval.max’ being set to 0.01 and ‘lfc.min’ to 0.1. Finally, a prism object containing all data required for running BayesPrism was created using the new.prism() function, and the deconvolution was performed using the run.prism() function. Two-sided Wilcoxon rank-sum test was performed to examine any statistically significant enrichment.

For the survival analysis, the CTCL bulk RNA-seq cohort was grouped into high and low abundance of B cells (both estimated by bulk deconvolution and mean expression of *CD79A* and *CD79B*) by the optimal cut point determined using the cutp() function in the survMisc R package. We performed multivariate analyses using the Cox proportional hazards model (coxph() function in the survival R package) to correct clinical covariates including age, gender, and tumour stage for the survival analysis. Kaplan-Meier survival curves were plotted to show differences in survival time using the ggsurvplot() function in the survminer R package.

### Inference of cell:cell interactions

We inferred potential cell-cell interactions using CellPhoneDB^65,66^ (version 4). Briefly, we randomly downsampled the CTCL object to 100 cells per fine-grained cell type per donor. The generated object was then used to run CellPhoneDB analysis with default parameters and thresholds. For the downstream visualisation, we used the R package ktplots (https://github.com/zktuong/ktplots). When filtering the inferred interactions between F2/F3, MoDC3 and tumour cells, we restricted ligand and receptor genes to DEGs from the analysis comparing fibroblasts derived from CTCL and the other three conditions (healthy skin, AD and psoriasis).

### Prediction of druggable targets using drug2cell

To predict potential druggable targets on B cells and malignant T cells, we ran drug2cell^46^ these two cell types together with benign T cells as a comparator. Drug2cell is druggable target prediction tool which integrates drug-target interactions from the ChEMBL database (https://www.ebi.ac.uk/chembl/) with single-cell data to comprehensively evaluate drug target expression in single cells. We first calculated per-cell scores of ChEMBL drug targets using d2c.score() function. Then, we performed differentially expressed analysis on ChEMBL drugs by comparing B cells, benign T cells and malignant T cells using scanpy tl.rank_genes_groups() function. When visualising the result, we separated malignant T cells by patients in order to show drugs that potentially function in multiple patients, given the strong inter-patient heterogeneity of CTCL tumours.

### Immunohistochemistry of FFPE samples for TOX and GTSF1

Immunohistochemical staining for TOX and GTSF1 was performed on skin samples from healthy skin, AD, psoriasis and CTCL. In addition to the skin samples collected for scRNA-seq, a further cohort of CTCL patients gave informed written consent for previous clinical samples to be used. Automated immunohistochemistry staining was performed by the Newcastle Molecular Pathology Node on formalin fixed paraffin slides using the Ventana Discovery Ultra autostainer (Roche) and the DISCOVERY ChromoMap DAB Kit. Antibodies used for staining were Anti-GTSF1 (HPA038877, Atlas Antibodies) and Anti-TOX (HPA018322, Atlas Antibodies). Scoring for TOX and GTSF1 was performed manually by an haematopathologist and dermatologist, reviewing the slides and deciding on an agreed approximation of positive staining. Identification of neoplastic T cells was based on their location, size and immunophenotype.

### Immunohistochemistry of FFPE samples for B cells and FDC

Immunohistochemistry staining was performed on FFPE skin samples of CTCL tumours. Skin biopsies were fixed in 4% formalin, then moved to 70% ethanol, dehydrated and embedded in paraffin. For tissues from Vienna and Newcastle cohorts, FFPE samples were cut into 4 μm sections, deparaffinized using a Neoclear (Sigma-Aldrich) and ethanol series and autoclaved in citrate buffer at pH 6.1 (Dako) to achieve antigen retrieval. Blocking with hydrogen peroxide was performed. Subsequently, slides tissue sections were subjected to automated immunohistochemistry staining (Autostainer, Dako Agilent) using anti-CD20 antibody (mouse monoclonal, clone L26, Dako M0755), anti-CD79a antibody (mouse monoclonal, clone JCB117, Dako M7050), or anti-CD21 antibody (mouse monoclonal, clone 1F8, Dako M0784), followed by visualisation (EnVision FLEX, Dako Omnis, Agilent). For analyses in the Stockholm cohort, FFPE samples were cut into 3.5 µm sections, deparaffinized and subjected to citrate buffer at pH 9 (Dako) to achieve antigen retrieval. Staining was manually performed using anti-CD20 antibody (clone EP459Y, Abcam ab78237) and secondary Goat Anti-Rabbit IgG H&L antibody (Abcam ab214880).

Stained sections were imaged and digitised with Scanscope CD2 (Aperio Technologies), Zeiss AxioScan.Z1 Slide Scanner and TissueFAXS scanning system (TissueGnostics). Image-based automated cell detection for all samples was performed with HistoQUEST software (TissueGnostics).

## Author contributions

M.H. and S.T. conceived and directed the study. E.F.M.P, N.R., P.B., C.J. and H.B. acquired patient samples. E.F.M.P, E.S., F.T., P.M. and E.P generated scRNA-seq and spatial transcriptomics datasets. J.S., N.Z., J.N., C.M.B. and R.C. performed immunohistochemistry. F.T. and N.C. performed RareCyte analysis. R.L. led bioinformatics analysis. J.S., H.B., J.N., R.C. acquired and interpreted images. R.L., J.S., M.H., S.T., E.F.M.P., B.O., W.T., H.G., A.F., C.M.B. analysed and interpreted the data. L.G., R.A.B., N.G., J.E., I.G., G.R., L.G., C.A., S.H., F.L., D.H. and P.M.B. interpreted the data. P.H., N.R., J.L. K.R., provided patient samples. J.S., R.L. and M.H. wrote the manuscript. R.L. and J.S. designed the manuscript figures. All authors read and edited the manuscript.

## Acknowledgements

We thank Aidan Maartens for critical reading of the manuscript. This study was funded by the Wellcome Human Cell Atlas Strategic Science Support (WT211276/Z/18/Z). The Wellcome Sanger Institute is supported by core funding from the Wellcome Trust (206194 and 108413/A/15/D). For the purpose of Open Access, the author has applied a CC BY public copyright licence to any Author Accepted Manuscript version arising from this submission. M.H. is funded by Wellcome (WT107931/Z/15/Z) and the NIHR Newcastle Biomedical Research Centre; J.S. is funded by the Clinician Scientist Fellowship of the Austrian Society for Dermatology (OeGDV). R.L. is funded by the British Heart Foundation. E.F.M.P. was funded by a Wellcome 4Ward-North Clinical Training Fellowship. N.J.R. is funded by the NIHR Newcastle Biomedical Research Centre, NIHR Newcastle In Vitro Diagnostics Co-operative and NIHR Newcastle Patient Safety Research Collaborative and is a NIHR Senior Investigator. H.B. is funded by the Swedish Society for Medical Research, the Swedish Cancer Foundation, Region Stockholm (clinical research appointment and ALF medicine), Welander and Radiumhemmet foundations. J.N. is supported by a Karolinska Institutet PhD student grant (KID). This publication is part of the Human Cell Atlas – www.humancellatlas.org/publications/[humancellatlas.org].

## Data and code availability

All raw sequencing data from this study have been deposited at EMBL-EBI ArrayExpress and are made publicly available at E-MTAB-12303. Our data can be explored on an online webportal, https://collections.cellatlas.io/ctcl. The code generated during this study is available at Github: https://github.com/ruoyan-li/Cutaneous-T-cell-lymphoma-study.

